# The *APOE* isoforms differentially shape the transcriptomic and epigenomic landscapes of human microglia in a xenotransplantation model of Alzheimer’s disease

**DOI:** 10.1101/2024.07.03.601874

**Authors:** Kitty B. Murphy, Di Hu, Leen Wolfs, Renzo Mancuso, Bart De Strooper, Sarah J. Marzi

## Abstract

Microglia play a key role in the response to amyloid beta in Alzheimer’s disease (AD). In this context, a major transcriptional response of microglia is the upregulation of *APOE*, the strongest late-onset AD risk gene. Of its three isoforms, *APOE2* is thought to be protective, while *APOE4* increases AD risk. We hypothesised that the isoforms functionally alter microglia by shaping their transcriptomic and chromatin landscapes. We used RNA- and ATAC-sequencing to profile gene expression and chromatin accessibility of human microglia isolated from a xenotransplantation model of AD. We identified widespread transcriptomic and epigenomic differences which are dependent on *APOE* genotype, and are corroborated across the profiling assays. Our results indicate that impaired microglial proliferation, migration and immune responses may contribute to the increased risk for late-onset AD in *APOE4* carriers, while increased DNA-binding of the vitamin D receptor in *APOE2* microglia may contribute to the isoform’s protective role.

## Introduction

Microglia are key players implicated in the genetic susceptibility and progression of Alzheimer’s disease (AD). AD genetic risk predominantly falls within regulatory regions of the genome, including those marked by H3K27ac^1^, an epigenetic modification found at active enhancers and promoters^2^. H3K27ac is dysregulated in the brains of individuals with AD^3,4^, and microglial H3K27ac regions are strongly enriched for AD genetic risk^5,6^. Microglia have also been associated with AD risk in their open chromatin regions^7–10^, and at the level of their transcriptome^11–13^. Recent research using single-cell transcriptomics has highlighted that microglia occur in various distinct subtypes and activation states, which are anticipated to exhibit different epigenomic and transcriptomic responses dependent on their environmental niche. This variation is expected to give rise to different downstream effects on AD pathogenesis. Supporting this is the continued characterisation of different microglial phenotypes in AD, including those responsive to amyloid beta (Aβ) aggregates, a pathological hallmark of AD^14–16^.

In mouse models, the strongest transcriptional response of microglia to Aβ aggregates is the upregulation of the gene *APOE*^14,17,18^, which harbours the strongest genetic risk factor for late-onset AD. *APOE* is involved in regulating cholesterol and other lipid transport across cells^19^. In the periphery, it is produced by macrophages in the liver, while in the brain, it is primarily produced by astrocytes^19^. In humans, *APOE* has uniquely evolved into three different isoforms: *APOE2*, *APOE4*, and *APOE4*. In AD, *APOE2* is thought to be protective, while *APOE4* increases disease risk up to 12-fold in homozygous individuals of certain human populations^20^. Several studies have begun to explore the role of microglial *APOE* in AD, using different samples and methodologies. In mouse models of AD, *APOE4* carriers exhibit a higher abundance of microglia stress and inflammatory markers, a phenotype also observed in human tissue^15^. Furthermore, *APOE4* microglia are linked to dysregulated lipid metabolism^21,22^, which subsequently triggers neurotoxicity and tau phosphorylation^22^. Regarding immune responses, *APOE4* microglia induce the signalling of transforming growth factor-β (TGF-β), a multifunctional cytokine, thereby hindering the appropriate, neuroprotective microglial response to AD pathology^23^. Clearly, *APOE* plays a critical role in regulating microglia in response to AD pathology. Thus far, studies have predominantly focused on *APOE4*, but it is equally important to determine the role of *APOE2*, which may potentially be antagonistic. We hypothesised that different *APOE* isoforms would differentially regulate the microglial phenotype in response to Aβ pathology. However, the unique evolution of *APOE* in humans makes it difficult to faithfully recapitulate its effects on AD risk and pathogenesis in mouse models. As previously suggested, investigating the microglial response in human tissue is challenging due to technical limitations and inconsistencies in biological findings^24^.

To tackle these challenges, researchers have developed a human microglia xenotransplantation model^25,26^, in which iPSC-derived human microglia are xenografted into the brains of mice. Single-cell profiling of these microglia has identified known and novel amyloid-responsive states^27^. These microglial states were enriched for different subsets of AD genetic risk genes, highlighting that multiple microglial states are influenced by AD genetic susceptibility. One of the primary human-specific microglial responses was impaired in *APOE4* microglia^16^. In addition to demonstrating the usefulness of this model in disentangling the microglial response to Aβ pathology, this highlights the need to investigate the functional role of AD genetic risk factors in a cell type-specific manner.

Here, we used ATAC-seq and RNA-seq to profile human microglia expressing the different *APOE* isoforms, which were xenotransplanted into the *App^NL-G-F^* mouse model of AD. This enabled us to delineate the effects of the different *APOE* isoforms on the epigenomic and transcriptomic landscapes of microglia in AD.

## Results

We transplanted isogenic iPSC-derived human microglia *APOE2*/0, *APOE4*/0, *APOE4*/0 and an *APOE* knockout (*APOE*-KO) into the brains of the *App^NL-G-F^* mouse model of Alzheimer’s disease^28^. At 12 months, by which point Aβ pathology is extensive^18,28^, microglia were isolated by FACS using human microglia-specific antibodies (CD11b+ hCD45+, **Fig. 1a**). This approach results in a scenario where the manipulations of the *APOE* genotype are restricted to microglia, thereby allowing us to study microglia-autonomous effects of the isoforms. To characterise the epigenomic and transcriptomic landscapes of these microglia, they were profiled using ATAC-seq for open chromatin and RNA-seq for gene expression, respectively. After quality control and pre-processing, we obtained high-quality chromatin accessibility data across 16 mice (*APOE2* = 5, *APOE3* = 5, *APOE4* = 4, *APOE*-KO = 2, **Supplementary Table 1**; **Supplementary Fig. 1**) and high-quality transcriptomic data across 17 mice (*APOE2* = 5, *APOE3* = 4, *APOE4* = 5, *APOE*-KO = 3, **Supplementary Table 1**; **Supplementary Fig. 1**). Overall, we observed widespread differences in gene expression and chromatin accessibility across microglia of the different *APOE* isoforms, highlighting the complexity of the microglial response to Aβ pathology. In support of the opposing roles of *APOE2* and *APOE4* in AD risk, the greatest differences were observed between these isoforms.

**Figure 1:**
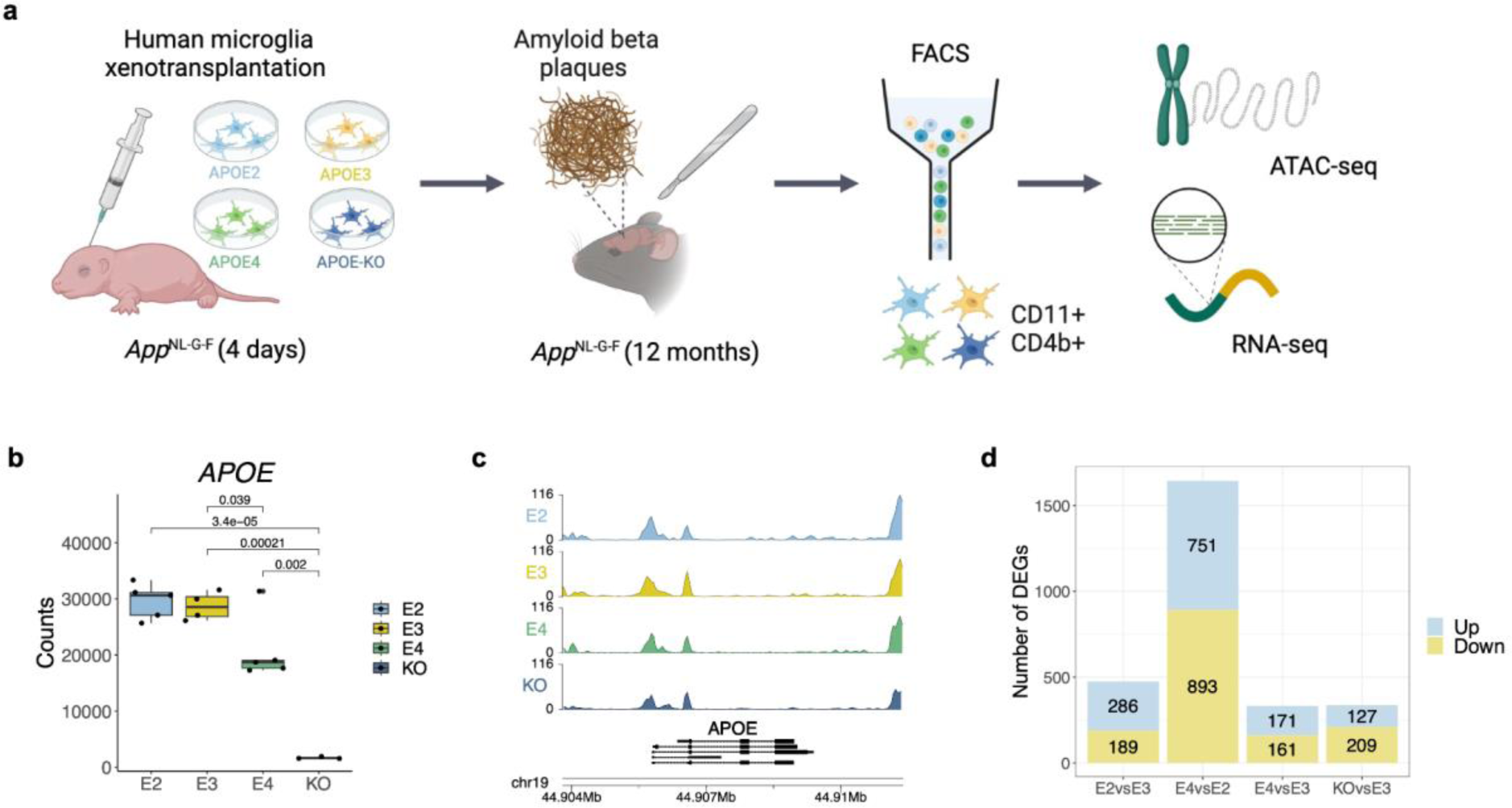
Transcriptomic and epigenomic profiling of xenotransplanted microglia reveals changes to their regulation in Alzheimer’s disease across the different *APOE* isoforms. **a** Experimental design for xenotransplantation of iPSC-derived human microglia into the brains of *App^NL-G-F^* mice (*APOE2* = 5, *APOE3* = 5, *APOE4* = 4, *APOE*-KO = 2) and high-quality transcriptomic data across 17 mice (*APOE2* = 5, *APOE3* = 4, *APOE4* = 5, *APOE*-KO = 3), followed by transcriptomic and chromatin accessibility profiling at 12 months. **b** Boxplot of expression profiles of *APOE*, confirming the knockout. **c** Genome tracks showing chromatin accessibility signals of all *APOE* groups around the *Apoe* locus. **d** Stacked barplot of the number of differentially expressed genes (DEGs; FDR < 0.05) identified through pairwise comparisons across the experimental groups.

### *APOE* isoforms are associated with consistent differences in gene expression and chromatin accessibility

*APOE* expression was significantly lower in three out of five knockout samples (**Supplementary Fig. 2**), and only these were used in downstream analyses (**Fig. 1b**). Additionally, *APOE* expression was lower in the *APOE4* microglia compared to *APOE2* and *APOE3*. This is in agreement with previous studies investigating the *APOE* variants in microglia and astrocytes^29,30^. Chromatin accessibility around the transcriptional start site (TSS) of *APOE* was consistent across the *APOE* groups (**Fig. 1c**). As the most commonly expressed allele^20^, we used *APOE3* as the baseline for pairwise comparisons in the differential expression and chromatin accessibility analyses performed using DESeq2^31^. In addition, we compared *APOE4* microglia with those expressing *APOE2*. When compared to *APOE3* microglia, differential expression analysis revealed 475 (286 up, 189 down, (FDR < 0.05); **Fig. 1d**, **Fig. 2a, Supplementary Table 2**) and 332 (171 up, 161 down, (FDR < 0.05); **Fig. 1d**, **Fig 2b, Supplementary Table 2**) differentially expressed genes (DEGs) in *APOE2* and *APOE4*, respectively. As expected, given their postulated opposing roles in AD risk, the direct comparison of *APOE4* with *APOE2* revealed the most differences, with 1,644 DEGs (751 up, 893 down, (FDR < 0.05; **Fig. 1d**, **Fig. 2c, Supplementary Table 2**). 127 genes were upregulated and 209 were downregulated in the *APOE*-KO when compared with the *APOE3* isoform (**Supplementary Fig. 3, Supplementary Table 2**).

**Figure 2:**
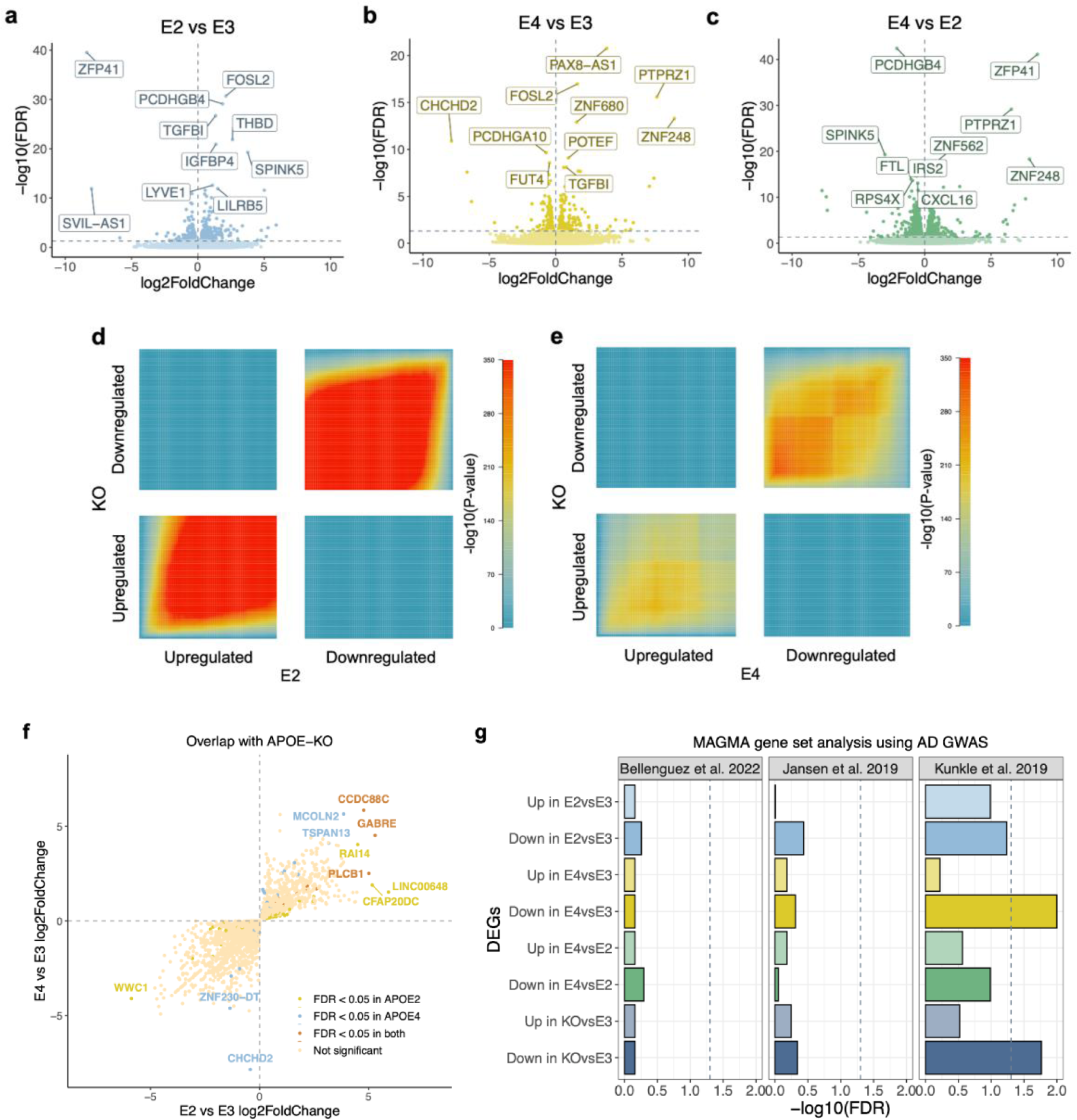
Microglia exhibit widespread differences in gene regulation across the *APOE* isoforms. **a** Differentially expressed genes in *APOE2* vs *APOE3* microglia. **b** Differentially expressed genes in *APOE4* vs *APOE3* microglia. **c** Differentially expressed genes in *APOE4* vs *APOE2* microglia. **d** Rank-Rank Hypergeometric Overlap (RRHO) heatmap comparing expression signatures between *APOE*-KO vs *APOE3* and *APOE2* vs *APOE3*. **e** RRHO heatmap comparing expression signatures between *APOE*-KO vs *APOE3* and *APOE4* vs *APOE3*. **f** Scatterplot of *APOE4* vs *APOE3* logFC against *APOE2* vs *APOE3* logFC for genes with expression profiles overlapping with the *APOE*-KO. **g** MAGMA gene set analysis using the differentially expressed genes across the *APOE* groups with three independent AD GWAS^35,37,38^.

Comparison with the *APOE*-KO enabled us to infer whether the transcriptional mechanisms underlying *APOE2* and *APOE4* microglia can be explained by loss and/or gain of function. We used Rank-Rank Hypergeometric Overlap (RRHO) to quantify the degree of overlap between expression signatures in the *APOE*-KO vs *APOE3* and *APOE2* vs *APOE3*, and the *APOE*-KO vs *APOE3* and *APOE4* vs *APOE3*. We observed significant overlap between expression patterns in the *APOE*-KO and *APOE2* comparison (Spearman’s rank correlation, rho = 0.55, p < 2.2e^-16^; **Fig. 2d**), as well as for the *APOE*-KO and *APOE4* variants (Spearman’s rank correlation, rho = 0.34, *p* < 2.2e^-16^; **Fig. 2e**). The strongest overlap was seen for genes downregulated in both *APOE2* and *APOE4*, suggesting that the mechanisms underlying the different *APOE* isoforms can be in part explained by a loss of *APOE* function. However, there were also unique transcriptional changes occurring in *APOE2* and *APOE4* microglia (**Fig. 2d-e**). This corroborates findings reported by Machlovi et al.^32^ in which the authors performed similar analyses investigating the *APOE4* and *APOE3* isoforms in mouse microglia. To further explore the overlaps with the *APOE*-KO, we correlated the genes that had similar expression profiles in *APOE2* and the *APOE*-KO, and *APOE4* and the *APOE*-KO, when compared to the *APOE3* isoform. All genes that had the same direction of expression change in *APOE2* and the *APOE*-KO, also had the same direction of expression change in *APOE4*, when compared with *APOE3* (**Fig. 2f**). A few genes were only significant in either *APOE2* or *APOE4*, including *TSPAN13*, which was upregulated in APOE4 microglia and is associated with lipid accumulation in microglia^22,33^. Due to the pleiotropic nature of *APOE*, we evaluated whether genes differentially expressed across the *APOE* groups exhibited differential enrichment for AD genetic risk variants. Using MAGMA gene set analysis^34^, we found that genes downregulated in the *APOE4* and *APOE*-KO microglia were enriched for risk variants identified in one AD GWAS^35^ (FDR < 0.05; **Fig. 2g, Supplementary Table 4**). This supports previous evidence suggesting that the mechanisms of *APOE4* reflect a loss-of-function in the context of AD^16,23,36^, with *APOE4* affecting biological pathways that are consistent with those linked to the polygenic component of AD.

To evaluate the upstream regulatory mechanisms associated with the transcriptomic changes across the APOE isoforms, we next investigated changes in chromatin accessibility. When compared to *APOE3* microglia, *APOE2* microglia had 40 differentially accessible regions (DARs) (24 up, 16 down, FDR < 0.05; **Fig. 3a, Supplementary Table 3**), and *APOE4* microglia had 50 DARs (38 up, 12 down, FDR < 0.05; **Fig. 3b, Supplementary Table 3**). Again, the direct comparison of *APOE4* with *APOE2* revealed the most differences, with 72 DARs (52 up, 20 down, FDR < 0.05; **Supplementary Fig. 4a, Supplementary Table 3**), with the fewest changes observed in the KO (13 up, 7 down, FDR < 0.05; **Supplementary Fig. 4b**). Notably, our analysis revealed consistent epigenomic and transcriptomic responses across microglia the different *APOE* isoforms (**Supplementary Fig. 5**). For instance, *CHCHD2,* a mitochondrial gene involved in promoting cellular migration^39^ and implicated in Parkinson’s disease (PD)^40,41^, was significantly downregulated in *APOE4* microglia when compared to both *APOE3* (logFC = –7.8, *p* = 1.3e^-11^; **Fig. 2b**, **Fig. 3c, Supplementary Table 2**) and *APOE2* (logFC = –7.4, *p* = 7.8e^-11^; **Fig. 2c**, **Fig. 3c, Supplementary Table 2**). In parallel, chromatin accessibility was significantly reduced close to the TSS of this gene when compared to *APOE3* (logFC = −7.8, *p* = 1.2e^-7^, distance to TSS = 0; **Fig. 3b, d, Supplementary Table 3**) and *APOE2* (logFC = −6.5, *p* = 8.3e^-9^, distance to TSS = 0; **Supplementary Fig. 4a, Supplementary Table 3**). Expression of this gene was also significantly reduced in the *APOE*-KO, suggesting a potential loss of protective function via this gene in the *APOE4* microglia (**Supplementary Fig. 3**). Similarly, the zinc finger protein *ZNF248* was upregulated in *APOE4* microglia (E4 vs E3, logFC = 9, p = 5.1e^-14^; E4 vs E2, logFC = 7.9, p = 5.2e^-19^; **Supplementary Table 2**) and genomic regions in the vicinity of this gene had increased chromatin accessibility (E4 vs E3, logFC = 6.3, p = 2.3e^-5^; E4 vs E2, logFC = 6.6, p = 3.2e^-5^; **Supplementary Table 3**). Interestingly, in an *in vitro* study investigating functional and transcriptional phenotypes of a TREM2 mutant and knockout in iPSC-derived microglia-like cells, *ZNF248* was upregulated in the TREM2-KO, while *CHCHD2* expression was reduced in the R47H mutant^42^. For an overall assessment of the concordance between ATAC-seq and RNA-seq, we correlated the logFC values between DEGs and DARs at the corresponding promoter peaks. We observed a strong correlation across all *APOE* comparisons, indicating general concordance between changes in chromatin and gene expression (**Supplementary Fig. 6**).

**Figure 3:**
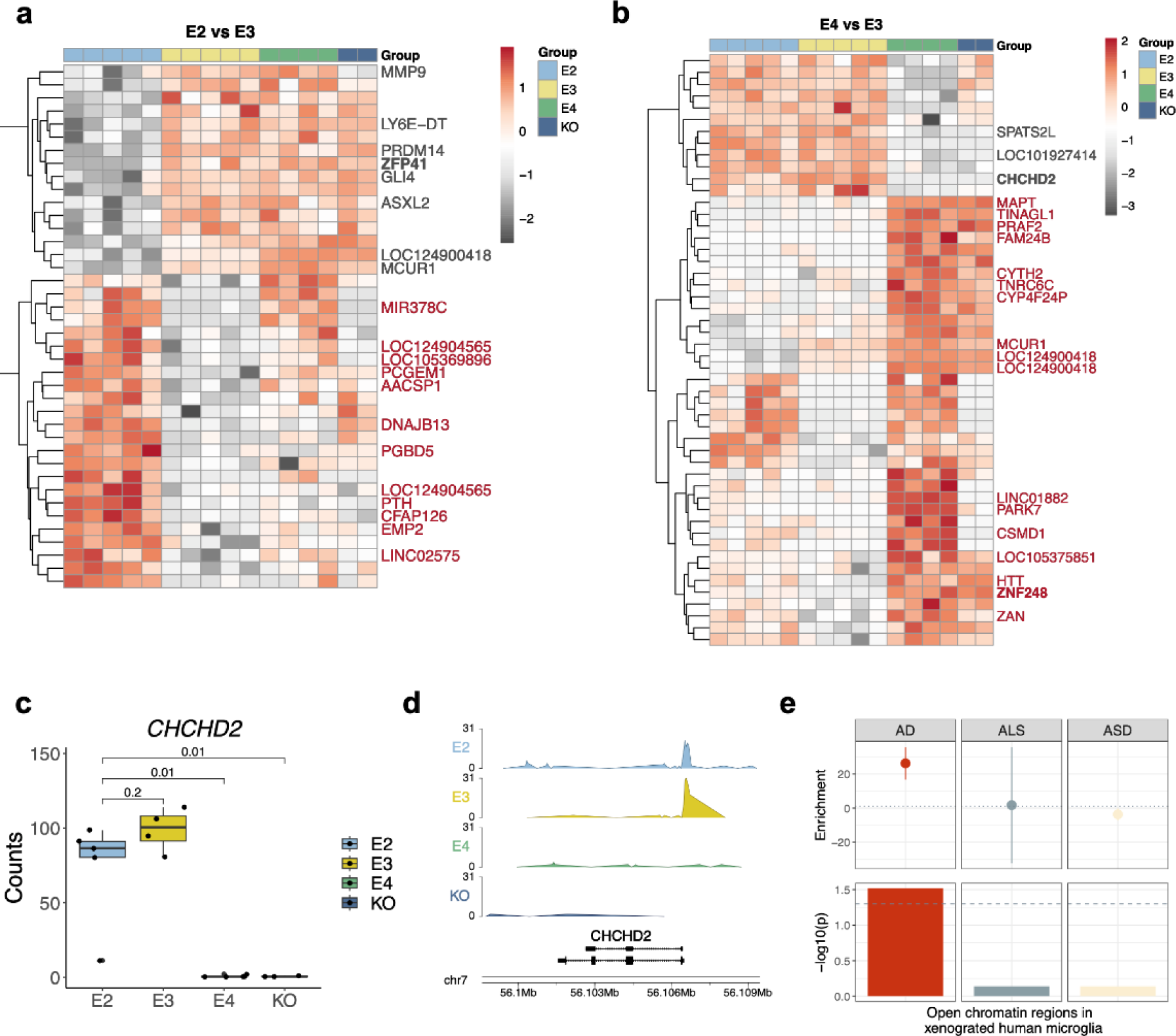
Concordant *APOE*-associated gene expression and chromatin accessibility signatures. Heatmaps showing differential chromatin accessibility of significant peaks (FDR < 0.05) when comparing **a** E2 vs E3, and **b** E4 vs E3. Shown are the genes annotated to the top 20 most significant peaks. Genes marked in bold were also significantly differentially expressed in the RNA-seq analysis. **c** Boxplot of expression profiles of *CHCHD2* shows reduced expression in the *APOE4* and the *APOE*-KO microglia. **d** Genome tracks of chromatin accessibility signals around the *CHCHD2* locus show a loss of the open chromatin peak at the *CHCHD2* promoter in *APOE4* and KO. **e** s-LDSC analysis using all open chromatin regions from the xenotransplanted human microglia with GWAS summary statistics for AD, ALS, and ASD shows a microglia-specific enrichment for AD.

Microglia-specific regulatory regions originating from human samples are strongly enriched for AD genetic risk^5,6,10,43^. To evaluate whether human iPSC-derived microglia xenotransplanted into the mouse brain would recapitulate this enrichment we used stratified linkage disequilibrium score regression (s-LDSC)^44^ with AD GWAS^37^. We found that open chromatin regions from the xenotransplanted microglia were enriched for AD heritability (FDR < 0.05, **Fig. 3e, Supplementary Table 5**). As the xenotransplanted microglia are predominantly responding to Aβ in our model, this enrichment also suggests that a significant proportion of AD risk is associated with microglial reactions to this pathological hallmark^45^. By repeating the analysis using GWAS data for autism spectrum disorder^46^, and amyotrophic lateral sclerosis (ALS)^47^, we confirmed that this enrichment was specific to AD, and not a general brain disease enrichment (**Fig. 3e, Supplementary Table 5**).

### *APOE2* and *APOE4* microglia show differential expression of cytokines

Human microglia from the xenotransplantation model used here were previously profiled using single-cell RNA sequencing^27^. Mancuso et al. (2024) report eight microglial states responsive to Aβ pathology, including previously characterised disease-associated microglia (DAM), as well as novel states annotated as cytokine response (CRM) and antigen-presenting response (HLA) microglia. Using hypergeometric testing, we found that genes dysregulated across the microglia the different *APOE* isoforms were strongly enriched within several microglia clusters (**Fig. 4a**): The strongest association was observed for genes downregulated in *APOE4* microglia, which were enriched in the HLA, ribosomal microglia (RM), and DAM clusters (**Fig. 4a**). Furthermore, genes downregulated in *APOE4* microglia and enriched in the DAM cluster were associated with negative regulation of tumor necrosis factor (TNF) cytokine production (**Fig. 4b**). Since negative regulation of cytokine production refers to processes that inhibit cytokine production, the downregulation of these genes in *APOE4* microglia suggests increased cytokine production in this isoform. CRM microglia mount a pro-inflammatory response driven by the upregulation of chemokines and cytokines, and have only been characterised in humans^16^. Furthermore, in response to Aβ, APOE*4* microglia shift to the CRM state rather than HLA^16^. Although we do not directly observe an association between *APOE4* microglia and CRM, we find an upregulation of chemokines and cytokines in *APOE4* when compared to *APOE2* (*CCL4L2*, *CCL3*, *CCL3L1*; **Fig. 4c-e**, **Supplementary Table 2**), with the exception of *CXCL16*, which was most highly expressed in *APOE2* microglia (**Fig. 4f**). In agreement, Machlovi et al. (2022), report increased cytokine production in *APOE4* microglia. Conversely, where the authors found increased *TNFα* in *APOE4* microglia, TNF family members, including *TNFRSF25* and *TNFRSF21*, were upregulated in *APOE2* microglia in our study. Taken together with the decreased HLA and DAM response in *APOE4* microglia here, these findings suggest that *APOE4* microglia fail to transition towards protective microglial states, and instead switch to more pathological states such as CRM (**Fig. 4a**).

**Figure 4:**
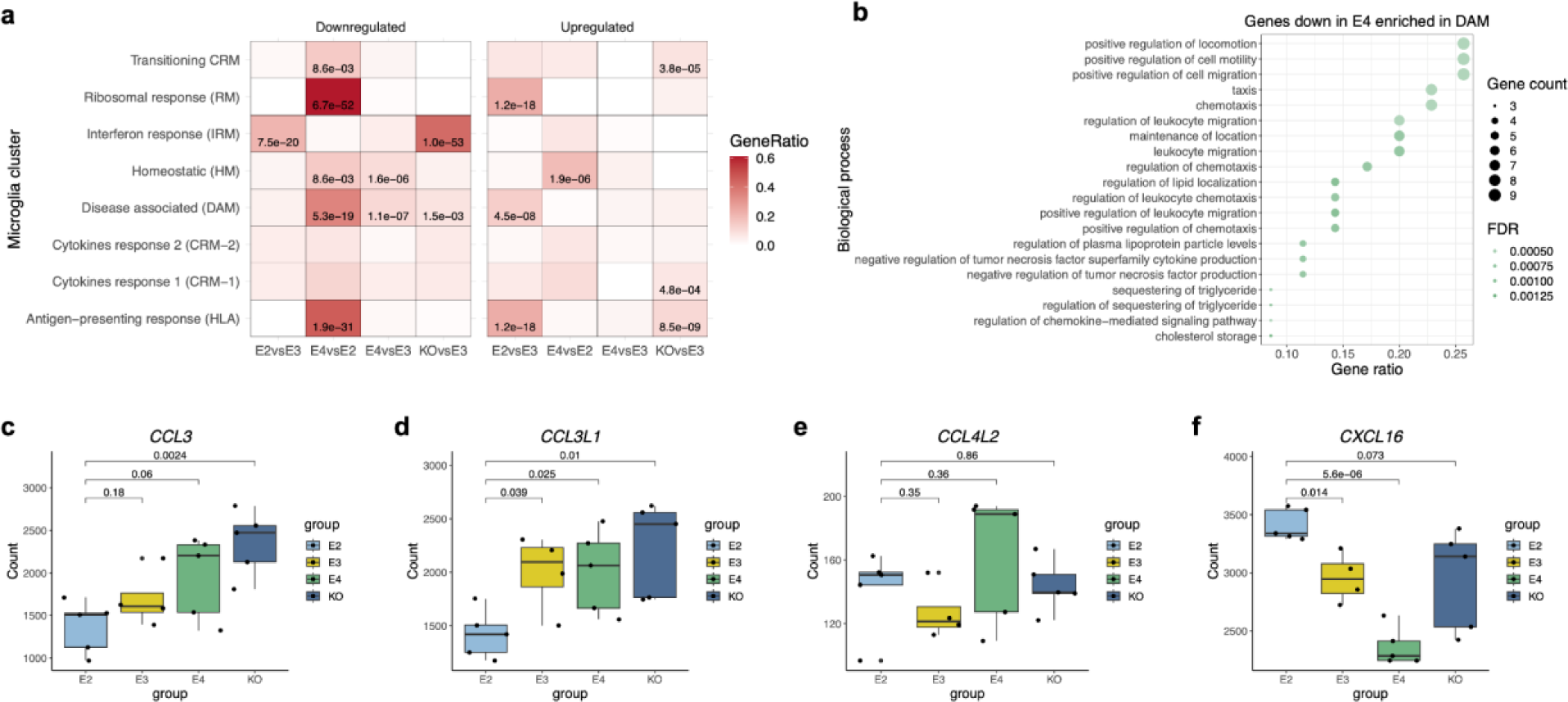
Pro-inflammatory cytokines are upregulated in *APOE4* microglia. **a** Heatmap showing enrichment of genes differentially expressed across the *APOE* groups amongst microglia clusters defined by scRNA-seq. **b** Dotplot of pathway enrichment analysis using genes downregulated in *APOE4* microglia that are enriched in the DAM cluster. **c-f** Boxplots of gene expression profiles of cytokines **c** *CCL3*, **d** *CCL3L1*, **e** *CCL4L2*, and **f** *CXCL16*.

### Gene networks upregulated in *APOE2* microglia are associated with cellular migration and immune response

We used weighted gene co-expression network analysis (WGCNA)^48^ to identify microglial gene modules with coordinated expression profiles. These modules were then tested for differential expression across the *APOE* groups, and the differentially expressed modules were functionally characterised using pathway enrichment analysis. We identified two differentially expressed modules significantly associated with GO biological processes. First, a gene module upregulated in *APOE2* microglia when compared to both *APOE3* and *APOE4* was associated with proliferation and cellular migration pathways (**Fig. 5a-b, Supplementary Table 6**). Such pathways are likely important for microglia being recruited towards the site of Aβ pathology and initiating its clearance, which in some cases, requires APOE^49^. Second, a module upregulated in *APOE2* when compared to *APOE4* was significantly associated with a range of immune responses, including both innate immune responses such as complement activation, and adaptive immune responses such as antibody-mediated immunity (**Fig. 5c-d, Supplementary Table 6**). Overall, the enrichment of proliferation, migration and immune responses, suggests enhanced protective microglial function in *APOE2*.

**Figure 5:**
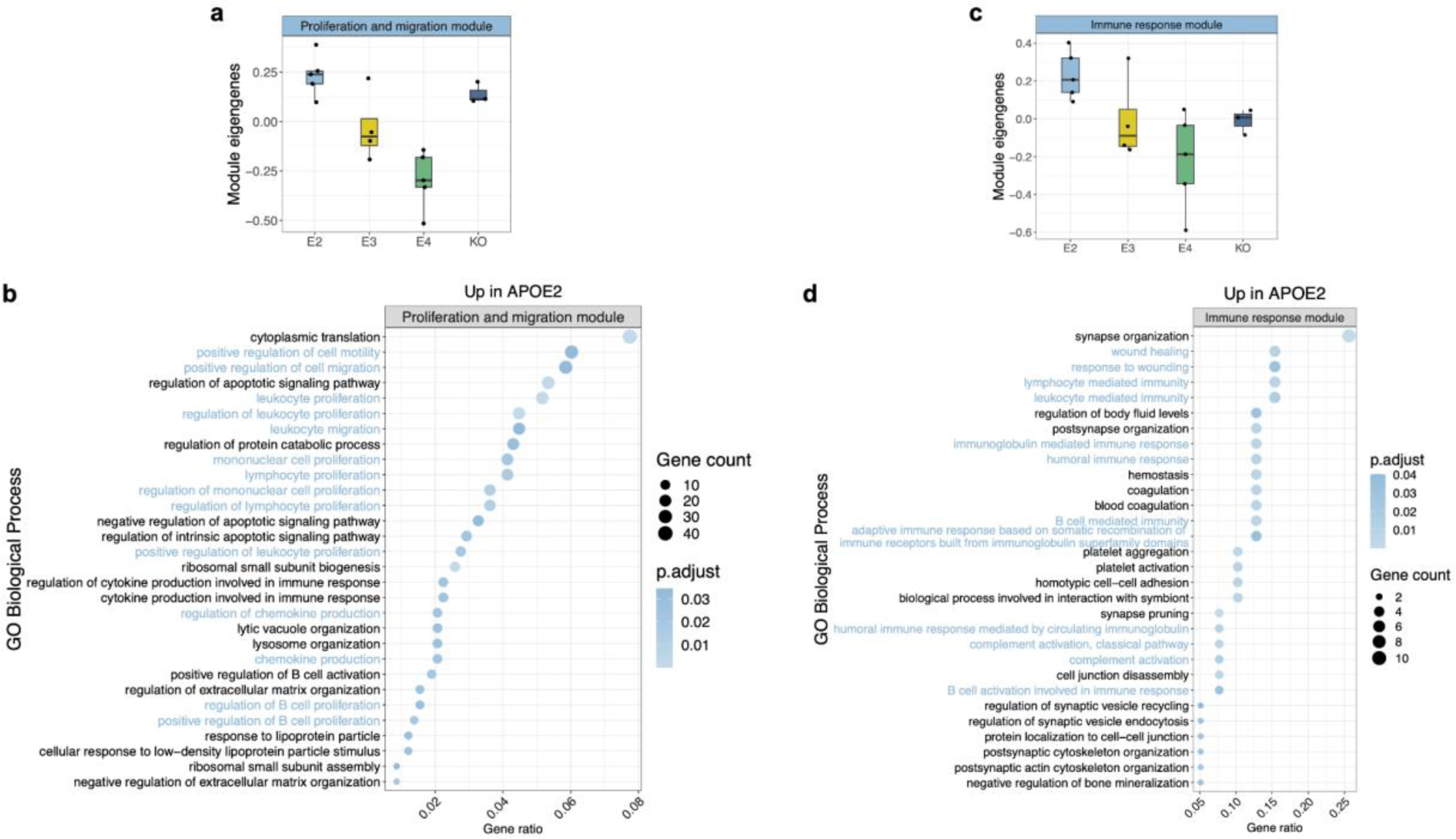
Gene networks upregulated in *APOE2* microglia are associated with cellular migration and immune responses. **a** Barplot of eigengene expression of the WGCNA module associated with proliferation and migration. **b** GO biological processes enriched for genes within the WGCNA module associated with proliferation and migration that is upregulated in *APOE2* microglia. **c** Barplot of eigengene expression of the WGCNA module associated with immune responses. **d** GO biological processes enriched for genes within the WGCNA module associated with immune responses, which is upregulated in *APOE2* microglia.

### Vitamin D receptor binding is upregulated in *APOE2* microglia

To better understand the regulatory machinery of human microglia in Alzheimer’s disease and the upstream orchestrators of altered transcriptional states, we performed de novo motif enrichment analysis using HOMER^50^ on the open chromatin regions from the xenotransplanted microglia. Specifically, we used the top 100 hyper- and hypo-acetylated peaks in *APOE2* and *APOE4* microglia as input and defined all ATAC-seq peaks as the background set. Regions with increased accessibility in the *APOE2* microglia were strongly enriched for the DNA binding motif of the vitamin D receptor (VDR) (**Fig. 6a**), a ligand-inducible transcription factor (TF) and main mediator of vitamin D signalling ^51^. Importantly, vitamin D deficiency has been linked to increased risk for AD^52,53^. We next assessed whether VDR target genes were upregulated in *APOE2* microglia. A hypergeometric test confirmed an overrepresentation of genes upregulated in *APOE2* when compared to both *APOE3* and *APOE4* (**Fig. 6b**), in a list of monocytic VDR target genes identified in a previous study^54^. This confirms the expected downstream transcriptional response predicted by increased VDR binding in *APOE2* microglia. Activation of an anti-inflammatory microglia phenotype via interleukin 10 (IL-10) and vitamin D signalling has been reported previously^55^. Consistent with this, in our data, the alpha subunit of the IL-10 receptor (*IL-10RA*) was significantly upregulated in *APOE2* microglia (**Fig. 6c**), suggesting increased anti-inflammatory signalling in the *APOE2* isoform.

**Figure 6:**
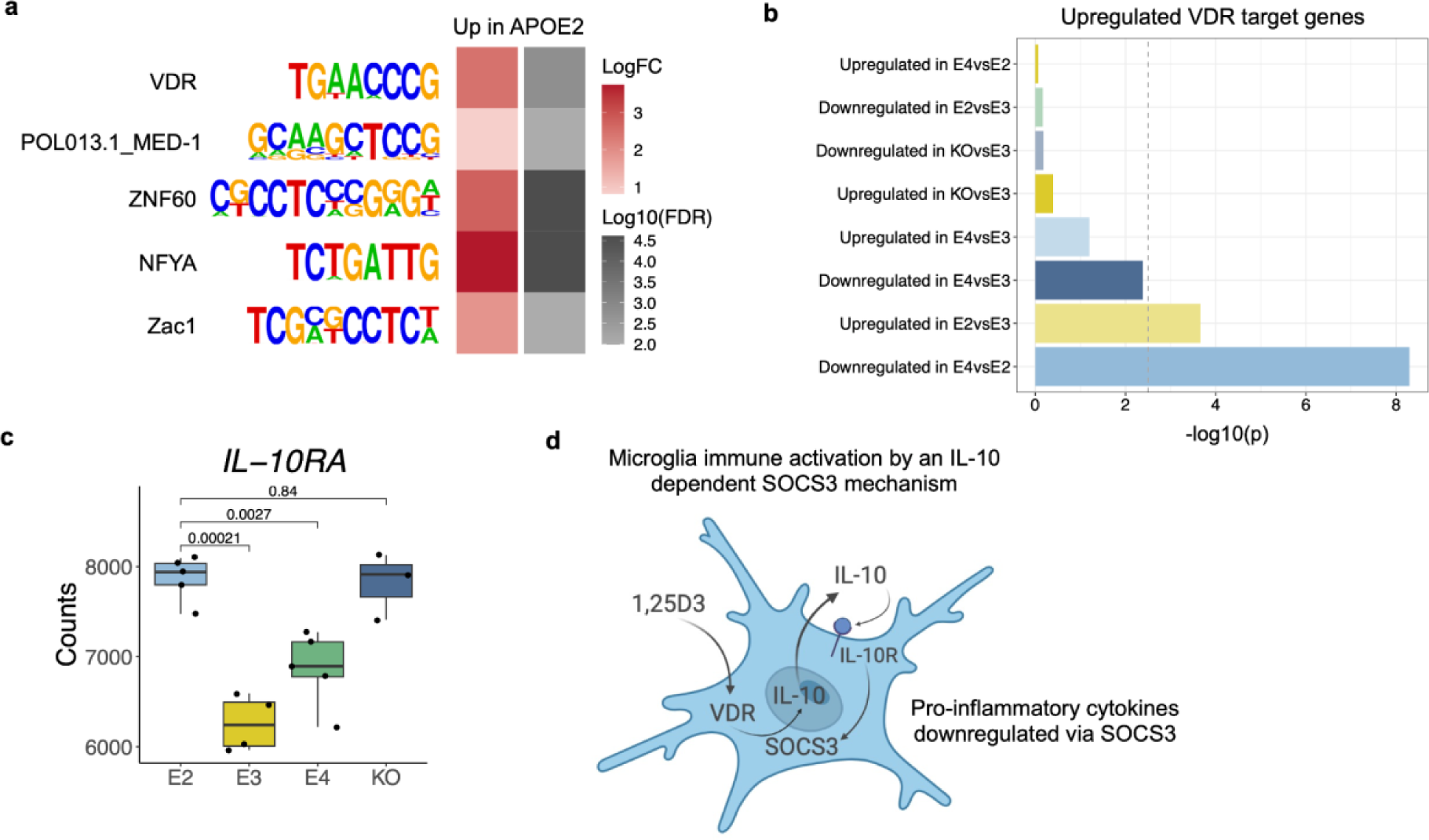
Regions with increased chromatin accessibility in *APOE2* microglia are enriched for the binding of vitamin D receptor. **a** Heatmap showing enrichment of motifs in regions with increased chromatin accessibility in *APOE2* microglia. **b** Barplot showing the overrepresentation of genes upregulated in *APOE2* microglia in a list of VDR target genes in monocytes^56^. **c** Boxplot of gene expression profiles of *IL-10RA*. **d** Graphic adapted from Boontanrart et al. (2016), showing a mechanism of anti-inflammatory microglia activation mediated through vitamin D and IL-10 signalling.

## Discussion

Increasing evidence points to a highly complex response of microglia to AD pathology. Here, show that the human *APOE2*, *APOE3*, and *APOE4*, differentially regulate microglia in the context of Aβ aggregates. By profiling human microglia isolated from a xenotransplantation model of late-onset AD using RNA-seq and ATAC-seq, we uncovered widespread changes to the transcriptomic and chromatin landscape of this cell type, dependent on the *APOE* isoform expressed. As anticipated, the largest differences were observed when comparing the AD risk opposing *APOE2* and *APOE4* microglia.

First, we observed consistent epigenomic and transcriptomic responses for several genes, including *CHCHD2* and *ZNF248*. *CHCHD2* is involved in promoting cell migration^39^ and has been linked to familial and sporadic Parkinson’s disease (PD)^40^, where it is transcriptionally downregulated. Its decreased expression and chromatin accessibility in *APOE4* microglia, but also in the knockout, suggest a potential loss of neuroprotective function in this isoform. Conversely, *ZNF248* was upregulated in *APOE4* and the knockout in both assays, suggesting a potential gain of toxic function. In a study comparing the effects of *TREM2* knockout and a *TREM2* mutation in a model of human microglia, the *TREM2* knockout had deficits in phagocytosis, chemotaxis, and survival that were not observed in the *TREM2* mutant^42^. *ZNF248* was one of only four differentially expressed genes with reduced expression in the knockout but increased expression in the mutant. Although the authors argue that it is unlikely that such a limited number of genes, including *ZNF248*, could explain such vast phenotypic differences^42^, the overlap between *APOE4* microglia and *TREM2* knockout microglia is interesting. The convergence of results between the DEGs and DARs highlights the robustness of using multiple independent assays to profile cellular states in a disease context. Overlapping expression signatures with the *APOE* knockout enabled us to infer whether the *APOE2* and *APOE4* isoforms resulted in a loss or gain-of-function. Aβ deposition is lower in AD mouse models with *APOE* knocked out^57–59^. Similarly, *APOE2* is associated with reduced Aβ pathology in both animal models and humans^60^. In contrast, the genes downregulated in *APOE4* microglia and the knockout were both enriched for AD genetic risk, lending support to previous reports of *APOE4* microglia increasing AD risk through loss-of-function mechanisms^16,23^. Further investigation into the genes shared between the knockout and *APOE4* microglia highlighted a strong upregulation of *TSPAN13*. This gene is also upregulated in microglia that accumulate damaging lipid droplets in the ageing brain ^33^, and in microglia homozygous for *APOE4*, in response to Aβ^22^. Our data suggest that lipid dysregulation in *APOE4* microglia may be driven by a loss of function.

Genes downregulated in *APOE4* were enriched within distinct microglial states identified in response to Aβ: HLA, RM, and DAM^27^. HLA represents a novel, human-specific microglial state that has a pronounced response to Aβ pathology and is thought to play a protective role^27^. RM are enriched for ribosomal genes. In murine AD models, stage 2 DAM cells, which signal the full activation of the DAM programme that is thought to be protective, are enriched for ribosomal genes^14^. When considered collectively, these enrichments suggest that *APOE4* microglia fail to shift into protective states. Furthermore, our analyses point toward diminished migratory capacity in *APOE4* and enhanced migratory capacity in *APOE2* microglia. Previously, *APOE4* microglia were shown to downregulate their expression of cellular migration genes in response to demyelination^61^, and pericytes derived from *APOE4* carriers exhibited downregulation of genes associated with cellular migration^62^. This also suggests that migratory capacity is not a cell type or pathology-specific mechanism affected by APOE in AD.

Another mechanism through which *APOE4* may be exerting its pathogenic role in AD is by mounting a pro-inflammatory response. In other studies, mouse microglia expressing the humanised *APOE4* allele increased cytokine production^32^, and xenotransplanted human microglia shifted towards a pro-inflammatory state^27^. Similarly, we report a general upregulation of cytokines in APOE4 when compared to both *APOE2* and *APOE3* microglia. One exception was *CXCL16*, which was upregulated in the APOE2 microglia. However, this chemokine has been reported to drive microglia to an anti-inflammatory phenotype in brain tumors^63^. Machlovi et al. (2022) reported increased *TNFα* expression in *APOE4* microglia, while we observed the opposite - TNF family genes were increased in *APOE2* when compared to *APOE4*. Therefore, it is important to consider that *APOE2* may also be driving a pro-inflammatory response, and while the isoform is thought to be protective in AD, it can increase risk for other neurological conditions^64^. This pro-inflammatory response may be balanced by the anti-inflammatory mechanisms suggested here and in other studies^65,66^. In addition, it remains unclear as to what triggers the increased production of pro-inflammatory cytokines in the *APOE* isoforms. For instance, in *APOE2* microglia the pro-inflammatory response may be triggered by their interaction and clearance of Aβ plaques. Whereas in *APOE4* microglia, the increased production of pro-inflammatory cytokines could be due to a lack of response to Aβ, which in turn would result in continued Aβ deposition and a sustained pro-inflammatory response.

Several studies have shown an enrichment of AD genetic risk within microglia-specific genes and regulatory regions from the human brain^5,6,11^. Here, using the chromatin accessibility profiles from the xenotransplanted microglia we recapitulate this enrichment, highlighting the robustness of this model for investigating human genetic risk in a disease context. At the level of the transcriptome, AD genetic risk was enriched within genes downregulated in *APOE4* microglia but also the *APOE* knockout. Supporting previous studies, this overlap suggests that AD risk increased by the presence of *APOE4* is partially mediated through loss of protective function. Our findings underscore the need to consider the interplay between genetic risk factors, and microglial states in AD.

In addition to arguing for increased proliferation, migration, and immune response in *APOE2* microglia as underlying this isoform’s protective effect, we report a potential upstream regulatory role for the VDR. In the context of AD, low levels of vitamin D have been associated with a higher incidence of the disease^52,53^, and vitamin D supplementation has been shown to improve disease outcomes^67,68^. It is important to take into consideration *APOE* genotype, as some studies have shown that *APOE4* carriers have higher vitamin D levels^69,70^ and therefore vitamin D supplementation may be more beneficial to non-carriers^68^. Vitamin D acts via binding to VDR, and enrichment of VDR in regions with increased chromatin accessibility in *APOE2* may therefore enable these microglia to be more responsive to vitamin D, regardless of serum levels. Furthermore, the increased expression of the *IL-10* receptor in *APOE2* microglia was particularly interesting. Vitamin D, via the VDR, increases the expression of the anti-inflammatory cytokine *IL-10*^55^. IL-10 then activates *SOCS3* via the IL-10 receptor, and this mechanism suppresses the expression of pro-inflammatory cytokines. Several other studies have also demonstrated an association between vitamin D and the expression of anti-inflammatory factors in microglia^71–76^. The functional role of VDR activation and binding warrants further studies in terms of mechanisms and therapeutic exploration.

Our study has several limitations. First, microglia exist in different subtypes and states. For example, microglia associated with Aβ plaques may have distinct transcriptomic and epigenomic profiles compared to less responsive microglia. While it is worth considering the contributions of *APOE* from mouse microglia and astrocytes, these should remain consistent across the *APOE* groups and, as such, should not affect differential expression and chromatin accessibility analyses. Although transcriptomic and epigenomic profiling provide valuable insights into gene regulatory mechanisms, other factors, such as histone modifications, also play a significant role in AD^3,4,43^. We linked regions with differential chromatin accessibility to differentially expressed genes based on their proximity. While this approach may capture promoter-gene relationships, many DARs may also function as enhancers. These enhancer-gene links can regulate target genes up to a megabase away, making them more challenging to identify. Using appropriate chromatin interaction data such as Hi-C could help disentangle these connections. Finally, while the absence of an adaptive immune system is necessary to prevent xenograft rejection, it may lead to unaccounted for changes in the microglial response^77^.

Our work sheds light on the regulation of microglia in AD: we show that it is dependent on *APOE* isoform, at both the level of the transcriptome and epigenome, further highlighting the complexity of this cell type in response to Aβ. Our work suggests that *APOE4* microglia have compromised microglial functions including diminished migratory capacity and heightened pro-inflammatory responses compared to *APOE2*, and these may underlie the increased risk of AD seen in carriers of this isoform. Furthermore, our findings underscore the importance of considering the interplay between genetic risk factors, such as *APOE*, and microglial states in disease progression. Importantly, we highlight the potential involvement of the VDR in modulating microglial responses, providing new avenues for therapeutic exploration. Overall, the use of the microglia xenotransplantation model coupled with genome-wide profiling has enabled us to dissect the regulatory landscape of microglia expressing the different APOE isoforms. In future, this approach could be extended to other relevant genes. In summary, our study emphasises the complex interplay between genetic, epigenetic, and environmental factors in shaping microglial responses in AD and underscores the need for targeted interventions based on *APOE* genotype.

## Methods

### Differentiation of microglial progenitors

Microglial progenitors were differentiated using the MIGRATE protocol, described in detail in Fattorelli et al. (2021)^26^.

### Human microglia xenotransplantation model

*App^NL-G-F^* mice were crossed with homozygous Rag2tm1.1Flv Csf1tm1(CSF1)Flv Il2rgtm1.1Flv Apptm3.1Tcs mice (Jacksons Lab, strain 017708) to generate the Rag2-/-Il2rγ-/-hCSF1KI *App^NL-G-F^*. In total, we transplanted 500,000 cells bilaterally across 20 mice. Mice had access to food and water ad libitum and were housed with a 14/10 h light-dark cycle at 21°C in groups of two to five animals. All experiments were conducted according to protocols approved by the local Ethical Committee of Laboratory Animals of the KU Leuven following country and European Union guidelines. Five biological replicates were prepared per experimental group: *APOE2*, *APOE3*, *APOE4*, *APOE* KO. From each sample, FACS purification of the following cell numbers were attained: 100,000 cells for ATAC-seq, 200,000 cells for RNA-seq. ATAC-seq samples were processed immediately after cell collection for tagmentation and elution of transposed DNA (details in ATAC-seq methods section). RNA-seq was conducted from cell pellets snap frozen in liquid nitrogen.

### RNA-seq library preparation

From snap frozen cell pellets of 200,000 cells per sample, RNA was extracted using the Monarch Total RNA Miniprep Kit (T2010) following the manufacturer’s instructions. RNA-seq was conducted using the rRNA depletion strategy rather than mRNA enrichment so that noncoding RNAs could be recovered^78^. rRNA depletion was performed using NEBNext rRNA Depletion Kit v2 Human/Mouse/Rat with RNA Sample Purification Beads (E7405), followed by stranded (directional) library preparation using the NEBNext Ultra II Directional RNA Library Prep Kit for Illumina (E7765) following manufacturer’s protocols without adjustments. RNA quality was checked using the Agilent RNA 6000 Pico Kit (5067-1513) and final libraries were assessed using Agilent High Sensitivity DNA Kit (5067-2646) where all libraries were appropriate for sequencing apart from one replicate of the *APOE3* microglia - therefore this isoform only has four biological replicates for RNA-seq.

### ATAC-seq library preparation

ATAC-seq was conducted as previously described^79^. Following FACS collection of 100,000 cells per sample, cells were spun down at 500*g* for 5 min at 4°C, and the supernatant was removed. Cell pellets were gently resuspended in 50 µL of ice-cold Lysis Buffer (10 mM Tris-HCl pH 7.4, 10 mM NaCl, 3 mM MgCl2, 0.1% IGEPAL CA-630). 2.5 µL Tagment DNA Enzyme (Illumina; 20034197) was added directly and gently mixed by pipetting. The transposition reaction was incubated at 37°C for 30 minutes, then transferred to ice. DNA was purified immediately with the Zymo ChIP DNA Clean and Concentration Kit (D5205) following manufacturer’s instructions. The DNA column was spun dry prior to elution of transposed DNA, which was conducted with 11 µL Elution Buffer. Purified DNA was stored at −20°C until library preparation. 10 µL DNA per sample was transferred into a PCR tube and 34.25 µL PCR master mix was added per sample. 6.25 µL of 10 µM Nextera Primer 2 (with barcode) was added per sample, where a different barcode was used for each sample to enable multiplexing. PCR was conducted using the following settings: (1) 72°C for 5 min, (2) 98°C for 30s; (3) 98°C for 10s; (4) 63°C for 30s; (5) 72°C for 1 min; (6) repeat steps (3)-(5) for a total of 10 cycles; (7) hold at 4°C. Amplified library was purified using the Zymo ChIP DNA Clean and Concentration Kit (D5205). The purified library was eluted using 20 µL Elution Buffer. 5 µL of 5x TBE Loading Buffer (Invitrogen; LC6678) was added and loaded in a 12-well 10% TBE gel (Invitrogen; EC62752BOX). A ladder was prepared using 0.25-0.5µL ORangeRuler 50 bp DNA Ladder (ThermoFisher; SM0613) diluted in 5 µL 5x TBE loading buffer. The gel was run at 70 V until DNA enters the gel, then increased to 140 V for approximately one hour. The gel was stained using 10 mL 1x TBE with SYBR Gold Nucleic Acid Gel stain (Invitrogen; S11494) diluted at 1:10,000 (1 µL). The gel was cut between 175-225 bp markers into a 0.5 mL DNA LoBind tube perforated three times with a 22G needle. The gel was shredded by centrifugation at maximum speed for 2 min at room temperature into a 1.5 mL DNA LoBind tube. 150 µL Diffusion Buffer (0.5 M Ammonium Acetate, 0.1% SDS, 1 mM EDTA, 10 mM Magnesium Acetate, ddH2O) was added to the gel in the 1.5 mL tube and shaken at room temperature for 45 minutes. The sample was then transferred to filter columns using wide-bore tips and spun at max speed for 2 min. DNA was purified (∼140 µL) using the Zymo ChIP DNA Clean and Concentration Kit and eluted with 10 µL Elution Buffer into 1.5 mL DNA LoBind tubes. Final libraries were quantified with the Qubit 1X dsDNA HS Assay Kit (ThermoFisher; Q33230) and stored at −20°C prior to sequencing (yield: ∼0.25 ng/µL).

### Sequencing

Final library size distributions were assessed by Agilent 2100 Bioanalyser and Agilent 4200 TapeStation for quality control before sequencing. Libraries were pooled to achieve an equal representation of the desired final library size range (equimolar pooling based on Bioanalyser/TapeSation signal in the 150bp to 800bp range). Paired-end Illumina sequencing using the HiSeq 4000 PE75 strategy was conducted on barcoded libraries at the Imperial Biomedical Research Centre (BRC) Genomics Facility following the manufacturer’s protocols.

### RNA-seq QC and processing

General QC of each sample was assessed using fastQC (https://www.bioinformatics.babraham.ac.uk/projects/fastqc/), followed by adapter trimming using TrimGalore! (https://github.com/FelixKrueger/TrimGalore). Reads were aligned to the GRCh38 genome and transcriptome using STAR^81^. Duplicate and multi-mapping reads were retained. Transcript quantification was performed using Salmon^82^, using the gc bias flag. Two of the *APOE* knockout samples with high *APOE* expression were excluded from subsequent analyses (**Supplementary Fig. 2**).

### ATAC-seq QC and processing

General QC of each sample was assessed using fastQC (https://www.bioinformatics.babraham.ac.uk/projects/fastqc/), followed by adapter trimming using TrimGalore! (https://github.com/FelixKrueger/TrimGalore). Reads were aligned to GRCh38 using bowtie2 with the following arguments: *–local –very-sensitive –no-mixed –no-discordant-I* 25 *-X* 1000. Post-alignment QC included removing: reads mapping to the mitochondrial genome, duplicate reads, multi-mapping reads, and reads with low mapping quality (q < 30). Read count generation was performed using featureCounts^80^. Additionally, peaks were filtered using the filterByExpr() function in DESeq2^31^, retaining only peaks with sufficiently high counts for statistical analysis. This left 228,041 peaks for downstream analyses. The 2 *APOE*-KO samples with high *APOE* expression in the RNA-seq data were also excluded from the ATAC-seq dataset. Additionally, another *APOE*-KO and one *APOE4* microglia sample were discarded because they did not meet QC standards.

### Differential expression and accessibility analysis

DeSeq2^31^ was used for the differential expression and differential chromatin accessibility analysis. DeSeq2 was designed for the differential analysis of RNA-seq data and has since been widely used for this purpose. In a recent study comparing methods for differential analysis of ATAC-seq read counts, Gontarz et al (2020) showed that with five replicates, which we have for most of our samples, DESeq2 had the lowest false positive rate and a true positive recall comparable to other methods available for differential accessibility analysis. For both analyses, the *APOE3* microglia samples were used as a baseline for comparison, and we additionally tested for differences between *APOE4* and *APOE2* microglia. To perform the differential analysis, we used the DESeq() function which provides a wrapper for three functions: estimateSizeFactors() for estimation of size factors, estimateDispersions() for estimation of dispersion, and nbinomWaldTest() for negative binomial GLM fitting and Wald statistics. Genes and peaks were defined as being significant if p < 0.05 after FDR correction.

### Weighted gene co-expression network analysis

To identify which genes had similar expression profiles across the *APOE* groups, we used weighted gene co-expression network analysis (WGCNA)^48^. First, transcripts with zero or low expression counts were filtered out using the filterByExpr() function in edgeR^83^. As suggested by the authors of WGCNA, the count data was normalised by variance stabilising transformation and explored for outliers using principal component analysis. An appropriate soft thresholding power was chosen to ensure a scale-free network and used as input to the blockwiseModules() function in WGCNA to calculate the adjacency matrix. This function was also used to detect gene co - expression modules and to calculate module eigengenes. As defined by the authors^48^, the module eigengenes are the first principal component of a given module and can be considered to represent the gene expression profile of that module.

### Functional enrichment analysis using differentially expressed WGCNA modules

The module eigengenes were used to perform differential expression analysis using lmFit() function in limma^84^, across the different *APOE* groups. The genes belonging to the top differentially expressed modules (FDR < 0.05) were then used as input to perform functional enrichment analysis using clusterProfiler^85^. As the only differentially expressed modules were associated with *APOE2*, the genes belonging to these modules were used to perform pathway enrichment analysis using clusterProfiler^85^, allowing characterisation of the *APOE2*-associated modules based on their gene ontology (GO) enrichments. GO terms were considered to be significantly associated with the given modules if p < 0.05 after FDR correction.

### Stratified linkage disequilibrium score regression

To estimate the proportion of disease SNP-heritability attributable to open chromatin regions in the xenotransplanted microglia, we performed stratified linkage disequilibrium score regression (s-LDSC). Annotation files were generated and used to compute LD scores. Publicly available GWAS summary statistics for a recent AD GWAS^37^ were downloaded and converted to the required format for LDSC. Steps for the analysis were followed as instructed here https://github.com/bulik/ldsc/wiki and required files were downloaded from https://alkesgroup.broadinstitute.org/LDSCORE/GRCh38. LDSC was run using the full baseline model, thereby computing the proportion of SNP-heritability associated with the annotation of interest, while taking into account all the annotations in the baseline model. As we observed a significant enrichment, we repeated the analysis using GWAS data for Amyotrophic Lateral Sclerosis^47^, and Autism spectrum disorder^86^, to ensure this was not a generic neurological enrichment.

### MAGMA gene set analysis

MAGMA gene set analysis^34^ was used to assess the enrichment of AD SNP-based heritability among differentially expressed genes across the *APOE* isoforms. The SNP window was restricted to the gene region (0,0). Summary statistics for three independent AD GWAS were downloaded^35,37,38^ and formatted for use with MAGMA using MungeSumstats^87^. P-values were corrected using the FDR method.

## Data and code availability

FASTQ and read count files have been deposited in the Gene Expression Omnibus (GEO) under accession GSE271384 for the ATAC-seq dataset and GSE271385 for the RNA-seq dataset. All the data and code required to reproduce the figures in this manuscript are available in our GitHub repository: https://github.com/neurogenomics/APOE_microglia. All supplementary tables are available at: 10.5281/zenodo.12516685

## Contributions

SJM, BDS and RM designed the study. DH, LW and RM performed experiments. KBM analyzed the results. KBM and SJM wrote the manuscript with input from all coauthors. SJM, BDS and RM supervised the project.

## Acknowledgements

We thank the members of the International Neuroimmune Consortium for their insightful conversations and support in this project. The scheme in Fig. 1a was created with BioRender.com

---

**Supplementary Figure 1:**
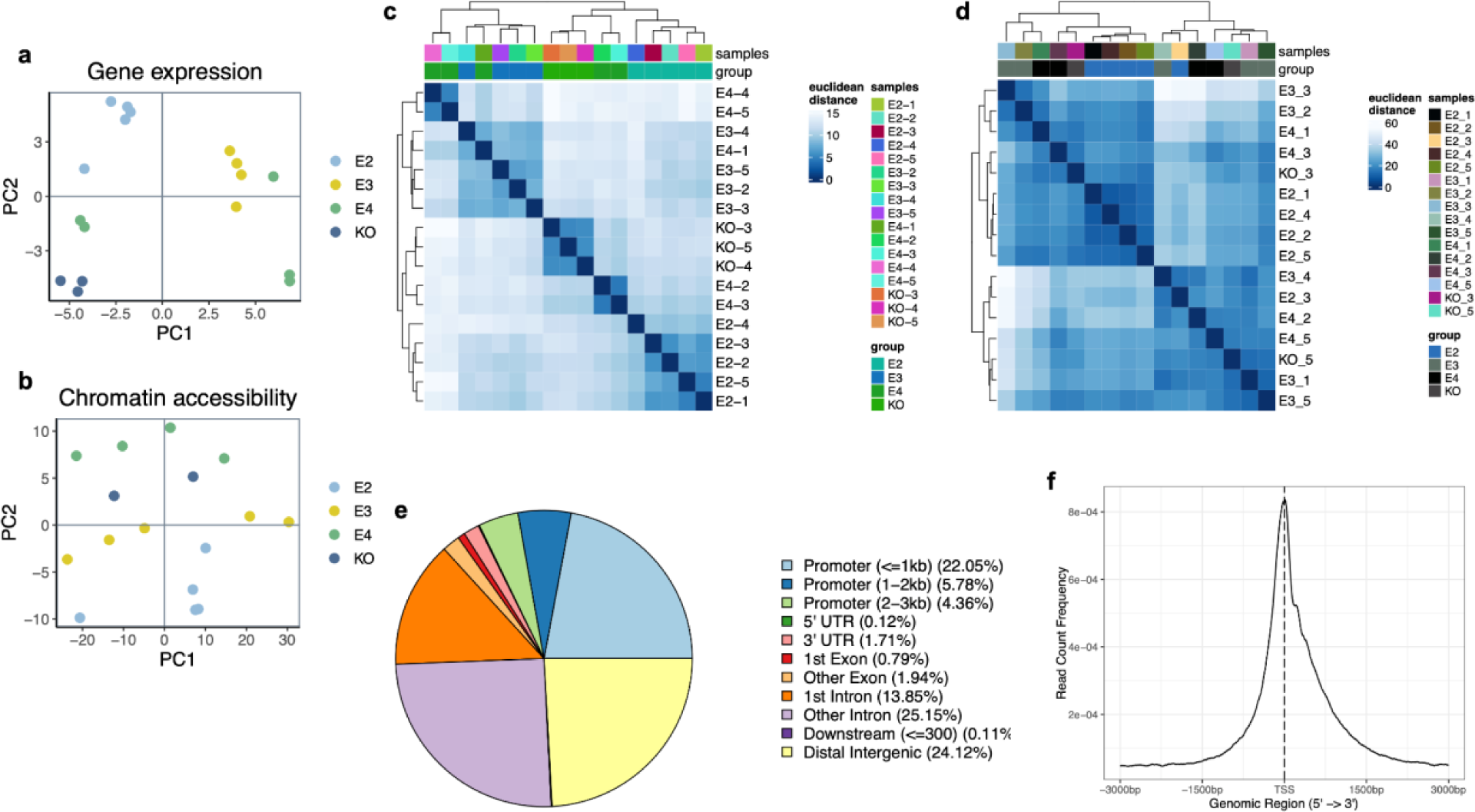
**a** PCA on gene expression across the *APOE* isoforms and the *APOE*-KO. PCA was generated using the variance stabilizing transformation (VST)-normalised expression counts matrix. **b** PCA on genome-wide chromatin accessibility across the *APOE* isoforms and the *APOE*-KO. PCA was generated using the variance stabilizing transformation (VST)-normalised expression counts matrix. **c** Hierarchical clustering using Euclidean distance of VST-normalised expression counts of the *APOE* isoforms and the *APOE*-KO. **d** Hierarchical clustering using Euclidean distance of VST-normalised ATAC-seq read counts of the *APOE* isoforms and the *APOE*-KO. **e** Pie chart of genomic annotations of the consensus set of chromatin accessibility peaks. **f** TSS enrichment profiles of ATAC-seq peaks.

**Supplementary Figure 2:**
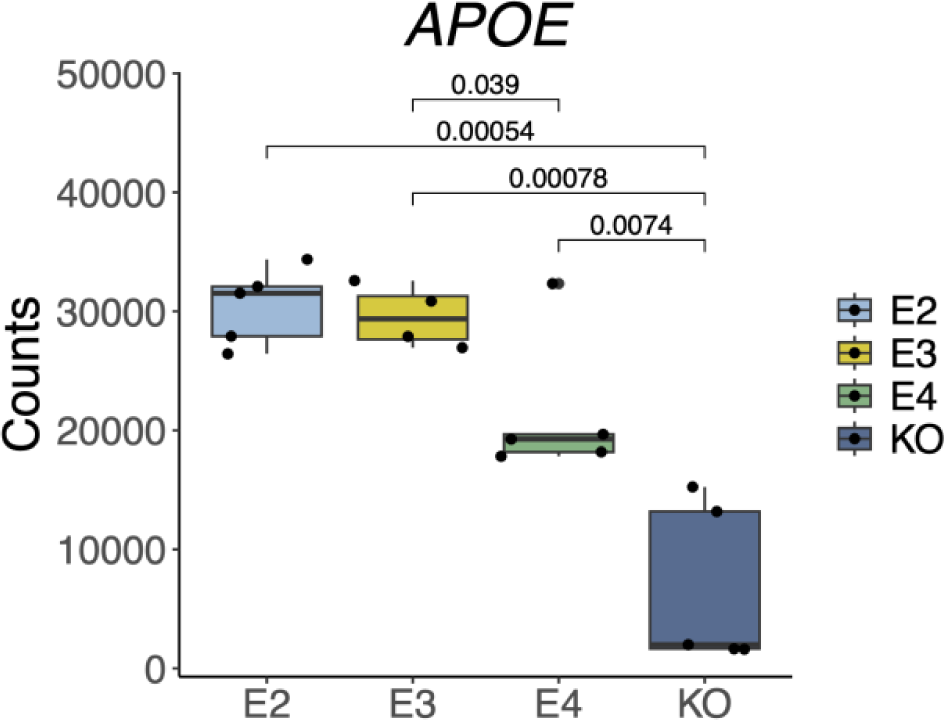
Boxplot of gene expression profiles of *APOE*, including two excluded *APOE*-KO samples which did not show loss of *APOE* expression.

**Supplementary Figure 3:**
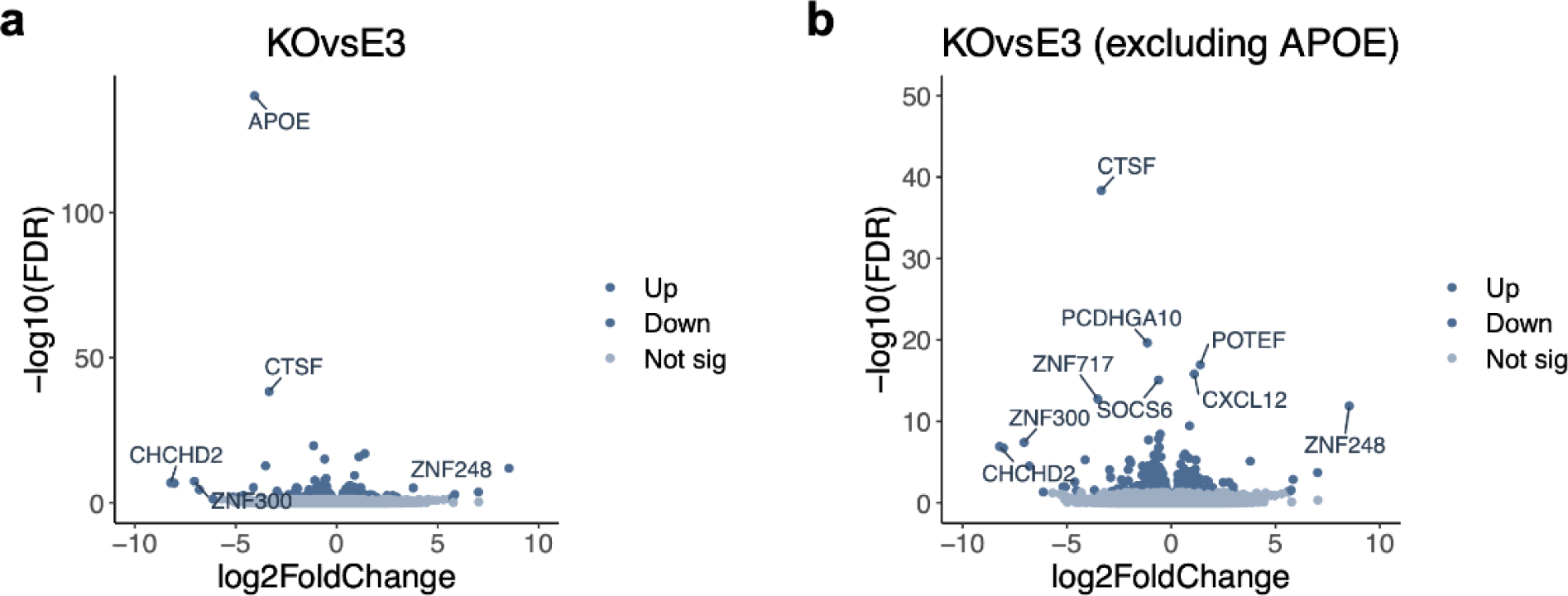
Differentially expressed genes in **a** *APOE* KO vs *APOE3*, **b** *APOE* KO vs *APOE3* (excluding *APOE*).

**Supplementary Figure 4:**
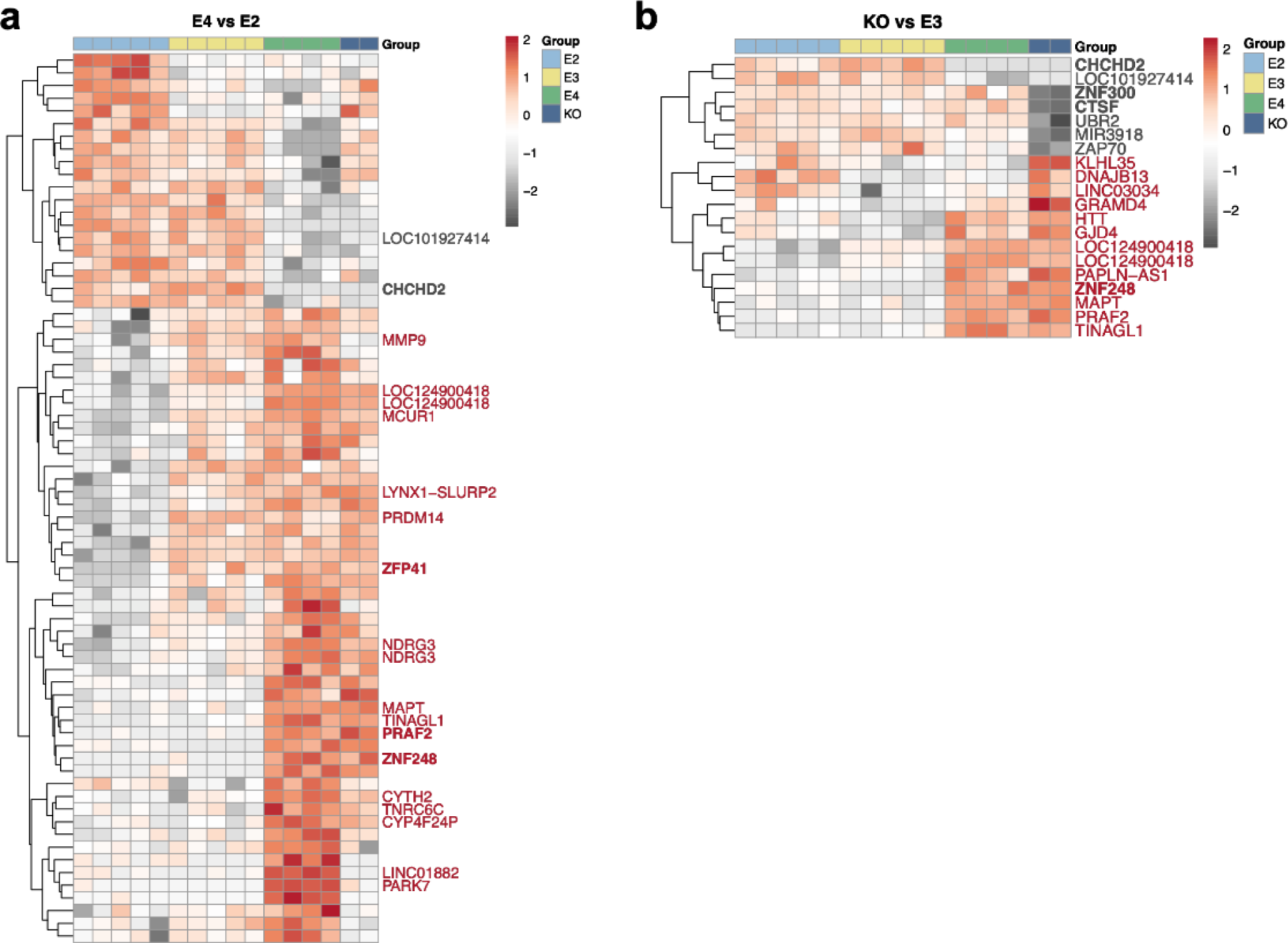
Heatmaps showing differential accessibility of significant peaks (FDR < 0.05) when comparing **a** E4 vs E2, **b** KO vs E3. Shown are the genes annotated to the top 20 peaks, genes marked in bold were also significantly differentially expressed in the RNA-seq analysis.

**Supplementary Figure 5:**
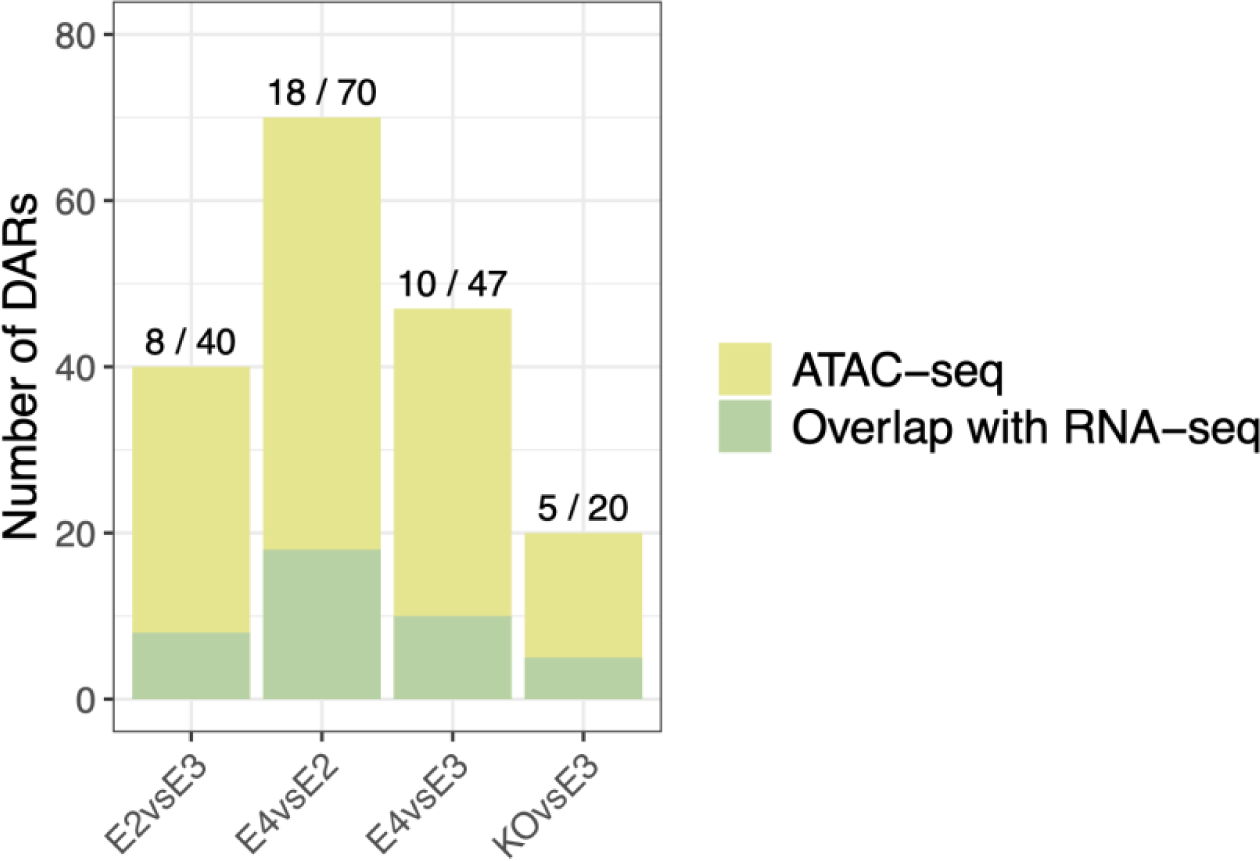
Stacked barplot of the number of differentially accessible regions (DARs) and how many of these overlap with the DEGs based on peak-to-gene annotation.

**Supplementary Figure 6:**
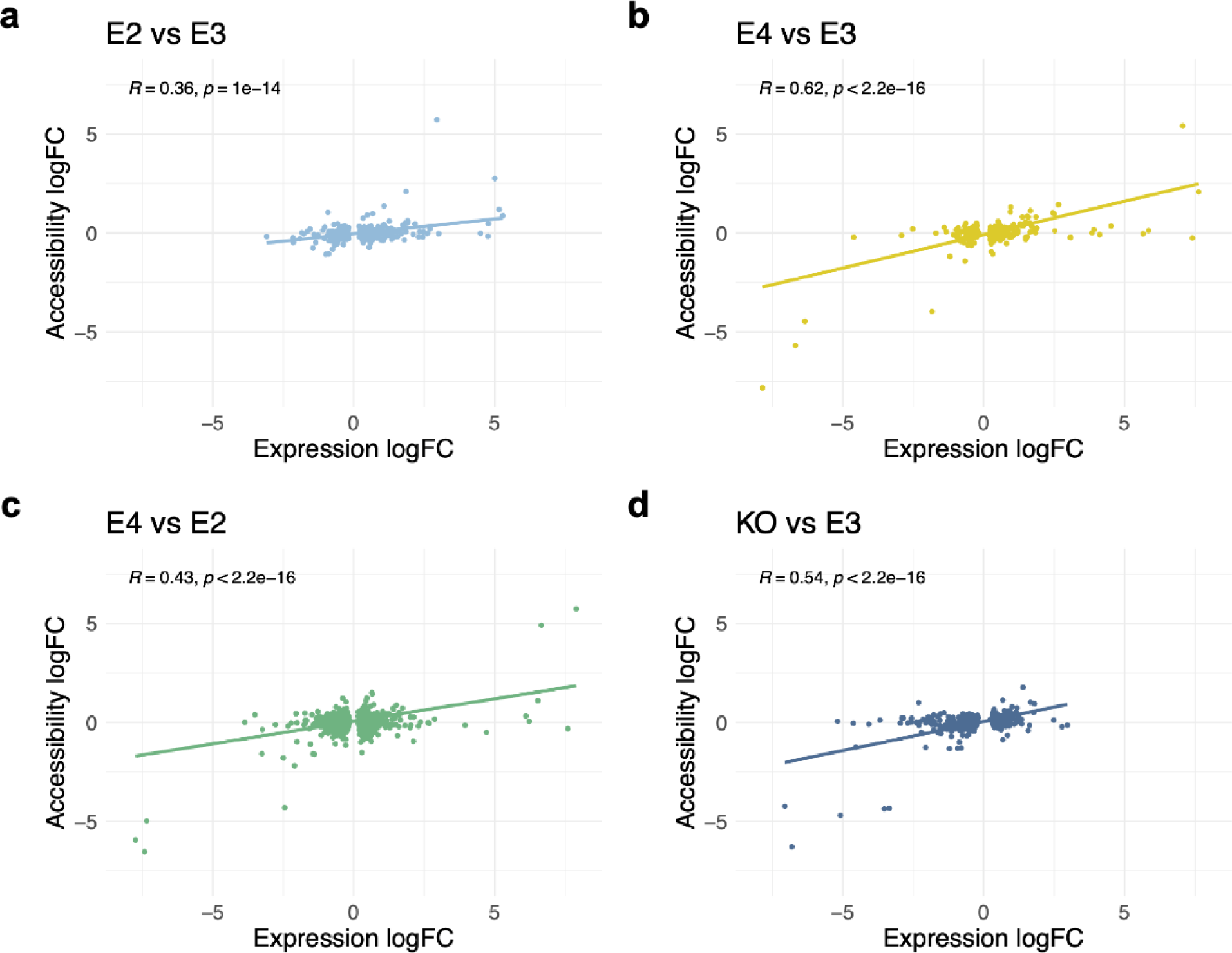
RNA-seq logFC and ATAC-seq logFC correlate across DEGs and their promoter peaks in **a** *APOE2* vs *APOE4*, **b** *APOE4* vs *APOE4*, **c** *APOE4* vs *APOE2*, **d** *APOE* KO vs *APOE4*. Correlations were performed using Pearson’s product moment correlation coefficient.

